# Sequential- *vs*. density gradient- centrifugation for the isolation of mitochondria-containing extracellular vesicles

**DOI:** 10.64898/2026.06.15.732469

**Authors:** Kandarp M. Dave, Bodhi T. Brady, Balaji Govindaswamy, Vivek S. Basudkar, Donna B. Stolz, Devika S Manickam

**Affiliations:** Graduate School of Pharmaceutical Sciences, Duquesne University, Pittsburgh, PA; Department of Biology and Microbiology, South Dakota State University, Brookings, SD; Department of Neurology, The University of Texas Health Science Center at Houston, TX; Center for Biologic Imaging, University of Pittsburgh Medical School, Pittsburgh, PA

**Keywords:** Extracellular vesicles, EVs, large EVs, mitochondria, brain endothelial cells, density gradient centrifugation, sequential centrifugation

## Abstract

A subset of extracellular vehicles (**EVs**) with particle diameters >200 nm, large vesicles (**lEVs**) contain mitochondria that increase recipient cell bioenergetics. To date, sequential centrifugation (**SC**) is the most reported protocol to separate lEVs from the smaller EVs (<200 nm)/exosomes. We have previously demonstrated that lEVs derived from brain endothelial cells (**BECs**) using the standard SC method transferred their innate mitochondria to recipient BECs, increased recipient BEC bioenergetics, reduced brain infarct volume, and improved behavioral outcomes in a mouse model of transient ischemic stroke. Despite their promising therapeutic activity, SC-isolated lEVs are likely a mixture of mitochondria-containing lEVs and non-mitochondria-containing lEVs. We hypothesized that subsequent purification of SC-isolated lEVs using density-gradient centrifugation (**DGC**) may yield a purer sample of mitochondria-containing lEVs. We established a DGC protocol to purify lEVs. In this pilot study, lEVs isolated using SC and DGC protocols were compared to determine their physicochemical characteristics and their effects on recipient BEC bioenergetics. SC-lEVs and DGC-lEVs both significantly restored ATP levels in OGD-injured BECs with no difference between groups. However, a Seahorse mitochondrial function assay revealed distinct functional effects: SC-lEVs did not significantly alter respiration, whereas DGC-lEVs induced a dose-dependent increase in oxygen consumption rate, indicating enhanced oxidative phosphorylation. These findings demonstrate that DGC purification yields a more mitochondria-enriched and functionally potent lEV preparation with an enhanced capacity to restore oxidative phosphorylation in ischemic BECs.

**Graphical abstract:** 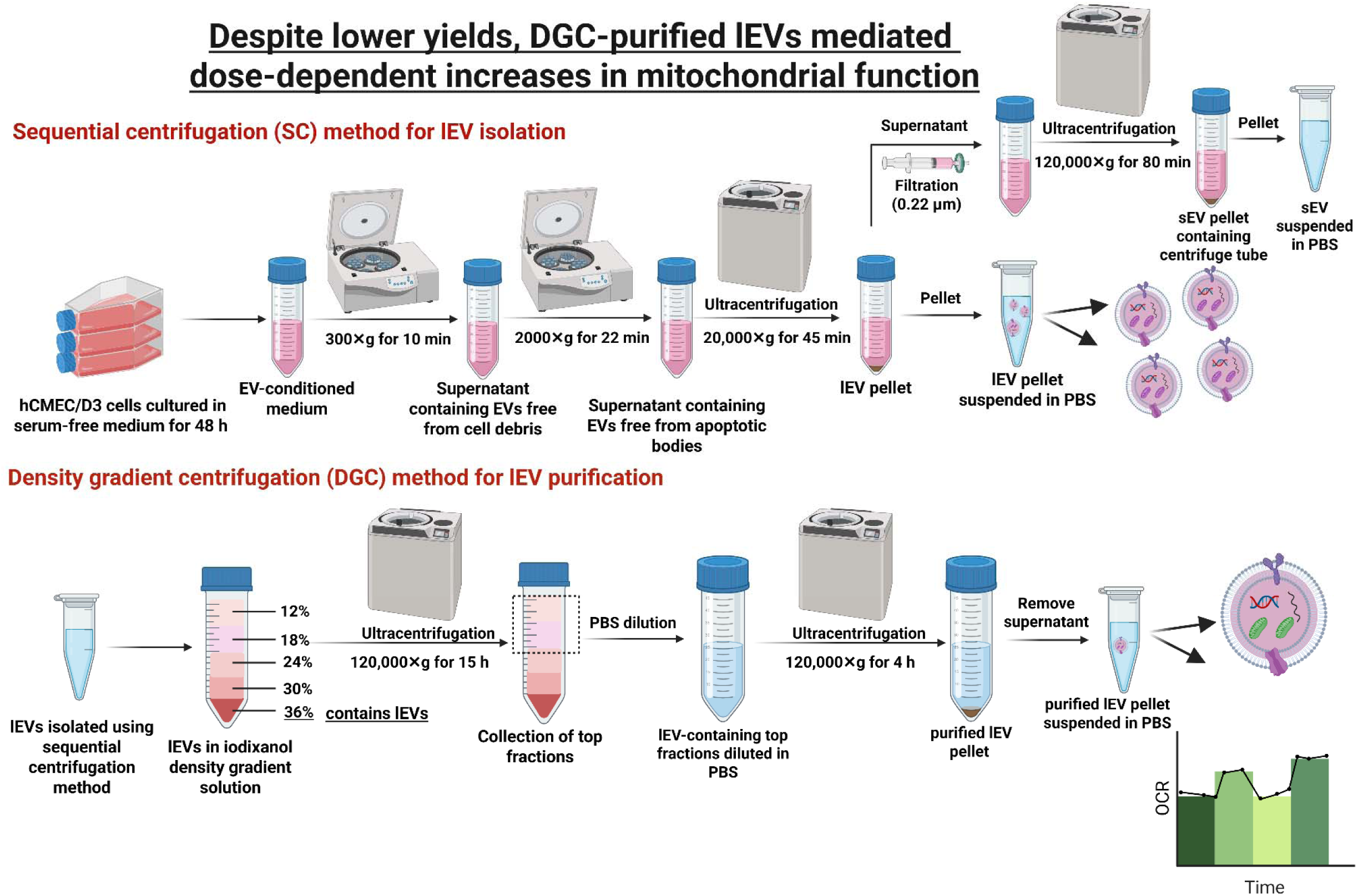

## 1. Introduction

Cell-derived extracellular vesicles (**EVs**) transfer RNA (1), DNA (2), lipids, and proteins (3) to recipient cells due to their known role in intercellular communication (4, 5). The low immunogenicity, tissue targetability, and ability to protect encapsulated biomolecules make EVs a suitable choice for drug delivery to the brain (4, 6, 7). Mitochondrial components containing EVs—including synaptosomes (8), exosomes, and large EVs (**lEVs**)—represent a promising strategy for mitochondrial delivery to target cells. Multiple studies including ours have confirmed the presence of mitochondria within lEVs (9–14) and their capacity to mediate mitochondrial transfer from donor to recipient cells (9, 12–15). Phinney *et al.* demonstrated that mesenchymal stem cells (**MSCs**) selectively package intact mitochondria into lEVs, while small EVs (**sEVs**) carry only mitochondrial DNA (16). lEV-mediated transfer of mitochondrial components to the injured cells has shown increased recipient cell ATP levels (17–20), oxidative phosphorylation (19), mitochondrial biogenesis, cell survival, and downregulation of proinflammatory cytokines (21) in various cell culture studies and preclinical disease models. Importantly, lEVs offer a natural vesicular platform for transferring functional exogenous mitochondria to target cells without the need for synthetic carriers such as polymer- or lipid-based nanoparticles.

lEVs are commonly isolated from biological fluids or culture-conditioned medium using sequential ultracentrifugation (**SC**) (22, 23), size-exclusion chromatography (24, 25), density gradient centrifugation (26), polymer-based EV precipitation (25), and affinity-based chromatography (27, 28). Among these, SC is the classical and most widely used approach, enabling the separation of different-sized EVs from large sample volumes through stepwise centrifugation at increasing speeds. This reagent-free process removes cells, debris, and apoptotic bodies while preserving EV morphology, surface properties, and functionality by avoiding cationic polymers, precipitating agents, or antibodies. Sequential centrifugation, therefore, remains a practical method for high-yield isolation of EVs in their native state, particularly when studying their intrinsic characteristics. However, lEVs isolated by SC (**SC-lEVs**) often co-precipitate with non-vesicular extracellular particles (26), free proteins, ribonucleoproteins, and lipoproteins (29).

Various methods can purify EVs from non-vesicular particles, protein aggregates, and lipoproteins, including size-exclusion chromatography, immunoaffinity capture, precipitation, and microfluidics (22, 30, 31). While effective, these approaches are often limited by low throughput, intermediate purity, dilution of EVs, cost, and accessibility. In contrast, density gradient centrifugation (**DGC**) provides superior purity and specificity while preserving EV functionality (31). DGC separates EVs from subpopulations and contaminants based on buoyant density, with sucrose and iodixanol (OptiPrep) being the most common gradient media (32). High-concentration sucrose gradients are hyperosmotic and viscous, potentially damaging vesicles and impairing resolution, and make it challenging to eliminate sucrose, which may interfere with downstream applications. Iodixanol, a non-ionic, water-soluble, biologically inert contrast agent (C_35_H_44_I_6_N_6_O_15_), forms isosmotic gradients up to 1.32 g/mL with low viscosity (32). The low viscosity of iodixanol gradients enhances particle penetration, improves separation resolution, and helps preserve the structural and functional integrity of EVs and mitochondria (31). Zhang *et al.* have developed high-resolution iodixanol density gradients that successfully separated exosomes from supermeres (33) and nonvesicular bodies (26). Brennen *et al.* showed that iodixanol DGC yields EVs with higher CD63 and minimal lipoprotein contamination compared to SC or size-exclusion chromatography (22).

Our previous work showed that SC-isolated lEVs (**SC-lEVs**) from BECs contain mitochondria and mitochondrial proteins and transfer them to recipient BECs under normoxic and hypoxic conditions (12–14). lEV treatment significantly enhanced ATP production and oxidative phosphorylation in hypoxic BECs. Importantly, these effects were abolished when donor BEC mitochondria were inhibited with rotenone or oligomycin prior to collecting EVs, confirming that the bioenergetic improvements depend on the innate lEV mitochondria (12–14). A single intravenous injection of SC-isolated lEVs into a mouse MCAo model reduced infarct volume and improved neurological outcomes (13, 14). These findings motivated us to investigate the feasibility of purifying lEVs obtained from SC isolation to further refine the effects of mitochondria-containing lEVs. In this study, we aimed to determine whether purifying SC-lEV mixtures could further enhance their mitochondrial and bioenergetic potency. In a pilot study, we established a DGC method to purify SC-lEVs. Human BEC-derived lEVs were first isolated via standard SC and then subjected to DGC using layered density solutions to separate lEVs from non-vesicular particles. Both SC- and DGC-lEVs were characterized for total EV protein content and physicochemical characteristics. We confirmed that SC-lEVs contain mitochondria via transmission electron microscopy, western blotting, and ImageStream analysis, and their capacity to transfer mitochondria to recipient BECs was evaluated. Finally, the effects of SC-and DGC-lEVs at different doses were compared to revive the ATP levels in recipient oxygen-glucose-deprived BECs. Our results indicated that SC-lEVs and DGC-lEVs both restored ATP levels in OGD-injured BECs with no clear difference between groups; however, only DGC-lEVs increased oxygen consumption rate in a dose-dependent manner, indicating enhanced oxidative phosphorylation/mitochondrial function.

## 2. Materials and Methods

### 2.1 Materials

| Material | Supplier |
| --- | --- |
| Endothelial cell basal medium | Lonza (Walkersville, MD) |
| Pierce Micro BCA and BCA protein assay kits | Thermo Scientific (Rockford, IL). |
| Collagen Type I | Corning (Discovery Labware Inc., Bedford, MA) |
| Hydrocortisone, human basic fibroblast growth factor, and ascorbic acid | Sigma-Aldrich (Saint Louis, MO) |
| Penicillin-Streptomycin solution, Chemically Defined Lipid Concentrate | Invitrogen (Carlsbad, CA) |
| Polycarbonate centrifuge tubes | Beckman Coulter, Inc. (Brea, CA) |
| OptiPrep Density Gradient Medium | Sigma-Aldrich (Saint Louis, MO) |
| MitoTracker Deep Red FM | Thermo Scientific (Rockford, IL) |
| Mouse monoclonal antibodies against ATP5A (cat#ab14748), GAPDH (cat#ab8245) | Abcam (Cambridge, MA) |
| Mouse monoclonal antibody against CD9 (cat#ab92726) | Life Technologies Corporation (Eugene, OR) |
| Alexa Fluor 790-conjugated donkey anti-mouse IgG | Jackson ImmunoResearch Lab Inc. (West Grove, PA) |
| Cell Titer Glo 2.0 reagent | Promega (Madison, WI) |
| Radioimmunoprecipitation assay buffer | Alfa Aesar (Ward Hill, MA) |
| Nitrocellulose membrane | GE Healthcare Life Sciences (Germany) |
| Intercept blocking buffer | LI-COR Inc. (Lincoln, NE) |

### 2.2. Cell lines and cell culture

A human cerebral microvascular brain endothelial cell line (**hCMEC/D3**, cat#102114.3C) was received from Cedarlane Laboratories (Burlington, Ontario) at P25 and used between P25 and P35 in all experiments. ***<u>We refer to hCMEC/D3 cells as BECs.</u>*** BECs were grown in tissue culture flasks and plates pre-coated using 0.15 mg/mL rat collagen I in a humidified 5% carbon dioxide incubator at 37 ± 0.5°C (Isotemp, Thermo Fisher Scientific). The cells were cultured in complete growth medium composed of endothelial cell basal medium supplemented with 5% fetal bovine serum, penicillin (100 units/mL)-streptomycin (100 μg/mL) mixture, hydrocortisone (1.4 µM), ascorbic acid (5 µg/mL), Chemically Defined Lipid Concentrate (1% v/v), 10 mM HEPES (pH 7.4), and human basic fibroblast growth factor (1 ng/mL). The complete growth medium was replenished every 48 hours until the cells formed confluent monolayers. Prior to passage, the cells were washed using 1x phosphate buffer saline (**PBS**) and detached from the flask using 1x TrypLE Express Enzyme (Gibco, Denmark). Cell suspensions stained with trypan blue (1:1 v/v ratio) were counted to calculate % live cells using a hemocytometer before passaging at 1:3 to 1:5 v/v ratios or plating in tissue culture plates at the indicated cell densities.

### 2.3. Isolation of large extracellular vesicles (lEVs) from BEC cell lines

#### 2.3.1. Isolation of lEVs using sequential centrifugation

lEVs were isolated from the conditioned medium of BECs using the differential ultracentrifugation method described earlier (34–36). Briefly, BECs were cultured in complete growth medium in tissue culture flasks with a growth area of 175 cm^2^ (T175). Confluent cells were washed with pre-warmed PBS pH 7.4 (0.0067M, PO_4_) without calcium and magnesium and incubated with serum-free medium for 48 h in a humidified incubator. Post-incubation, the EV-conditioned medium was collected in polypropylene tubes and centrifuged at 300 ×g for 11 min, followed by 2000 ×g for 22 min to pellet down cell debris and apoptotic bodies, respectively, using an Eppendorf 5810 R 15-amp centrifuge (Eppendorf, Germany). The supernatant was transferred to polycarbonate tubes and centrifuged at 20,000 ×g for 45 min at 4°C to pellet lEVs using a Sorvall MX 120+ micro-ultracentrifuge (Thermo Scientific, Santa Clara, CA). The supernatants were then discarded. The lEV pellets were washed with PBS twice, finally suspended in 300 μL of PBS, and stored at -80 °C until further use. lEVs isolated using SC are referred to as **SC-lEVs**.

#### 2.3.2. Purification of SC-isolated lEVs using density gradient centrifugation

The high-resolution iodixanol DGC-based EV purification protocol was adapted from Zhang *et al.* (26). First, lEVs were isolated from the conditioned medium of BECs using sequential centrifugation (**SC**), as described in ***section 2.3.1***. SC-isolated lEV pellets were suspended in PBS and subsequently pooled into a centrifuge tube. For DGC, about 700 μL of SC-isolated lEVs was transferred into a separate centrifuge tube. The remaining SC-lEVs were stored for comparisons with DGC-purified EVs. The stock solution of ice-cold 60% iodixanol was diluted with ice-cold PBS to prepare four different concentrations of iodixanol (30%, 24%, 18%, and 12%), as listed in **Table 1**. The crude pellet of SC-isolated lEVs was suspended in ice-cold PBS and mixed with ice-cold iodixanol for the final 36% (w/v) iodixanol solution (**Table 1**). lEVs containing 36% iodixanol solution were added to the bottom of a polycarbonate centrifuge tube. Solutions of descending concentrations of iodixanol (30%, 24%, 18%, and 12%) in ice-cold PBS were carefully layered on top, yielding a complete gradient. The bottom-loaded 12-36% (w/v) gradient containing centrifuge tubes was carefully transferred into a Sorvall MX 120+ micro-ultracentrifuge (Thermo Scientific, Santa Clara, CA) and subjected to ultracentrifugation at 120,000 ×g for 15 h at 4°C. Post-centrifugation, out of twelve individual fractions of 1 mL, the first seven fractions were pooled into a separate polycarbonate tube, whereas the remaining lower five mL fractions for independent characterization. These lower fractions were designated as non-vesicular extracellular particles (**NVEPs**). Both DGC-lEV and NVEP fractions were subsequently diluted with ice-cold PBS to a final volume of 20 mL and mixed well. The tube was then subjected to ultracentrifugation at 120,000 ×g for 4 h at 4°C. Post centrifugation, the supernatant was discarded, and the purified lEV pellet was suspended with 300 μL of PBS and stored at -80 °C until further use. lEVs isolated using DGC are referred to as **DGC-lEVs**, whereas the corresponding non-vesicular extracellular lEV particle fractions are referred to as **NVEP-lEVs**.

**Table 1:** Preparation of iodixanol (OptiPrep) gradients with ice-cold PBS. Crude lEVs were mixed with iodixanol to make a final iodixanol concentration of 36%. The blank 36% iodixanol solution was prepared with ice-cold PBS.

| **% of Iodixanol** | **Volume of 60% Iodixanol (μL)** | **Volume of PBS (μL)** | **Volume of Sample (μL)** |
| --- | --- | --- | --- |
| 36 – Sample | 1440 | 260 | 700 |
| 36 – Blank (no IEVs) | 1440 | 960 | 0 |
| 30 | 2500 | 2500 | 0 |
| 24 | 2000 | 3000 | 0 |
| 18 | 1500 | 3500 | 0 |
| 12 | 1000 | 4000 | 0 |

### 2.4. Total lEV protein measurements using micro bicinchoninic acid (BCA) assay

SC and DGC-lEVs were quantified by measuring their total protein content using a micro BCA protein assay for all studies described herein. For the micro BCA protein assay, 10 µL of lEVs suspended in PBS were lysed using 150 µL of 1*×*RIPA buffer. A 150 µL volume of the EV lysates or BSA standards (2-40 µg/mL) was pipetted into a 96-well plate, and an equal quantity of the micro BCA WR (reagent A: reagent B: reagent C = 25:24:1) was added to each well. After two hours of incubation at 37°C, the absorbance was measured at 562 nm using a SYNERGY HTX multi-mode reader (BioTek Inc., Winooski, VT).

### 2.5. Dynamic light scattering

Particle diameters and zeta potential of SC and DGC-lEVs were measured using dynamic light scattering. lEVs were diluted to a final concentration of 0.1 mg lEV protein/mL in either PBS pH 7.4 for particle diameters or in deionized water for zeta potential measurements. lEV samples were analyzed using a Malvern Zetasizer Pro (Malvern Panalytical Inc., Worcestershire, UK). All samples were analyzed in triplicate. Average particle diameter, dispersity index, and zeta potential values were reported as mean ± standard deviation (**SD**) of three independent experiments.

### 2.6. Western blot analysis of EV protein markers

SC-lEVs were evaluated for characteristic protein biomarkers using western blotting. BEC lysates were used as positive controls. BEC lysate (10 µg) and 50 µg lEV proteins were mixed with laemmli sodium dodecyl sulfate sample buffer and denatured at 95°C for 5 min. The samples were loaded onto a 4% stacking and 10% resolving polyacrylamide gel. Premixed molecular weight markers (Protein standards, ladder, 250 kDa-10 kDa, cat#1610374, BioRad Laboratories Inc.) were used as a reference control to confirm molecular masses of the protein bands in the sample lanes. The gel was run at 120 V for 120 min in Tris-Glycine pH 8.8 running buffer using a PowerPac Basic setup gel electrophoresis assembly (BioRad Laboratories Inc.). The proteins were transferred onto a 0.45 µm nitrocellulose membrane at 75 V for 90 min using a transfer assembly (BioRad Laboratories Inc.). The membrane was washed with 0.1% Tween 20 containing Tris-buffered saline pH 7.4 (T-TBS) and blocked with Intercept blocking buffer solution (LI-COR Inc., Lincoln, NE) for one hour at room temperature. The membrane was incubated with mouse anti-GAPDH (1 μg/mL), mouse anti-ATP5A (1 μg/mL), rabbit anti-TOMM20 (1 μg/mL), rabbit anti-calnexin (1 μg/mL), and rabbit anti-ARF-6 (1 μg/mL) primary antibodies in blocking buffer and T-TBS solution (1:1::v/v) overnight at 4°C. The membrane was washed with T-TBS and incubated with anti-mouse or anti-rabbit AF790 secondary antibodies (0.05 μg/mL) for one hour at room temperature. The membrane was washed with T-TBS and scanned using an Odyssey M imager (LI-COR Inc., Lincoln, NE) at 700 and 800 nm channels.

### 2.7. ImageStream analysis of calcein-AM and MitoTracker labeled EVs

#### 2.7.1. MitoTracker-based EV mitochondria and Calcein-acetoxymethyl (AM) ester-based EV staining

For labeling lEV mitochondria, lEVs were isolated from BECs prestained with Mitotracker deep red (**MitoT-red**). Confluent BECs in T175 flasks were stained with 100 nM MitoT-red in serum-free medium for 60 minutes in a humidified incubator. Post-incubation, the old medium was removed, and cells were washed with PBS pH 7.4. The cells were incubated in serum-free medium for 48 hours in a humidified incubator. Post-incubation, the conditioned medium was collected in polypropylene tubes. MitoT-red-lEVs were isolated using the differential ultracentrifugation method described in ***section 2.3.1*.** MitoT-red-sEVs were isolated from MitoTracker-stained BECs using sequential centrifugation to compare the relative abundance of MitoTracker-positive EVs between lEVs and sEVs. For calcein-AM-based EV labeling, 200 μL of MitoT-red-lEVs and MitoT-red-sEVs were incubated with 50 μM calcein-AM for 30 min in the dark on a mechanical orbital shaker. Post incubation, the samples were vortexed and covered with aluminum foil until further use for ImageStream analyses.

#### 2.7.2. ImageStream analysis of MitoTracker and calcein-AM dual-labeled EVs

ImageStream includes the combination of flow cytometry with fluorescence imaging technology that can be utilized for smaller particles such as EVs (37). About 100,000 total events of MitoT-red and calcein-labeled lEVs were captured in the SSC area versus SSC intensity plots, where calcein-positive lEVs (green fluorescence channels) were gated to separate all events from the calibration beads. A total of 5,000 calcein-positive events were acquired for further gating MitoTracker-positive events (red fluorescence) on red fluorescence channels. Single color stains (*i.e.,* only calcein+ve lEVs and only MitoTracker+ve lEVs) and calibration beads were used to adjust spectral compensations. Separate channels were used for side-scatter and brightfield images.

### 2.8. lEV mitochondria transfer into recipient BECs

#### 2.8.1. Isolation of mitochondria-labeled lEVs

For labeling lEV mitochondria, lEVs were isolated from BECs prestained with Mitotracker deep red (**MitoT-red**) as described in ***section 2.7.1*.** The total protein content in MitoT-red-lEVs was determined using a micro BCA assay.

#### 2.8.2. Uptake of MitoT-red-lEV into recipient BECs using fluorescence microscopy

lEV mitochondria transfer into recipient BECs under normoxic conditions was studied using fluorescence microscopy. Recipient BECs were cultured in 24-well tissue culture plates at 100,000 cells/well in complete growth medium for 72 hours. The cells were treated with MitoT-red-lEVs at 10, 30, and 60 μg EV protein/well in complete growth medium for 48 hours in a humidified incubator. Untreated cells incubated with complete growth medium were used as a control. Post-incubation, the treatment mixture was removed, and cells were washed with PBS. Cells were incubated with Hoechst stain (1 μg/mL) for 15 min. The cells were incubated in phenol red-free DMEM with 10% FBS medium prior to observation under an Olympus IX 73 epifluorescent inverted microscope (Olympus, Pittsburgh, PA) using Cyanine-5 (Cy5, excitation 651 nm and emission 670 nm) and DAPI channels at 20*×* magnification. Images were acquired and analyzed using CellSens Dimension software (Olympus, Pittsburgh, PA).

### 2.9. Effects of SC- and DGC-isolated lEV treatment on relative ATP levels in the oxygen glucose-deprived recipient BECs

The effect of SC-lEVs, DGC-lEVs, and NVEP-lEVs on the relative cellular ATP levels in the recipient BECs under OGD conditions was measured using a Cell Titer Glo-based ATP assay. BECs in complete growth medium were cultured at 16,500 cells per well in a 96-well plate. The cells were incubated in a humidified incubator at 37°C for 72 h. Post-incubation, the cells were washed with PBS pH 7.4 and treated with OGD medium for four hours in an OGD chamber pre-flushed with 5% carbon dioxide, 5% hydrogen, and 90% nitrogen at 37±0.5°C. Post-incubation, the OGD medium was removed, and cells were treated with SC- or DGC-lEVs diluted at different concentrations in the OGD medium for 24 h. Cells treated with OGD medium alone were used as the OGD control, and cells treated with complete growth medium were used as normoxic control. After incubation, the treatment mixture was removed, and the cells were washed and incubated with complete growth medium. An equal volume of Cell Titer Glo reagent was added. The plate was incubated for 15 min on a mechanical orbital shaker. A 60 µL aliquot of the mixture was transferred to a white opaque plate, and relative luminescence units at 1 s integration time were measured using a plate reader (Synergy HTX multimode plate reader, BioTek Inc., Winooski, VT). Relative ATP levels were calculated by normalizing the relative luminescence units (**RLU**) of treatment groups to the RLU of control, untreated cells (**Equation 1**).

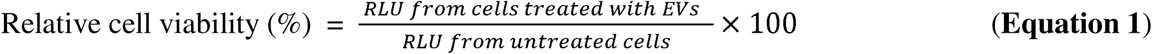

For EV and/or mitochondrial membrane disruption studies, SC-lEVs were treated with 0.1% Triton X-100 for 30 min to disrupt membrane integrity, followed by dilution in PBS and centrifugation at 20,000 ×g for 45 min. The resulting pellet was resuspended in PBS and evaluated for its ability to restore ATP levels in oxygen-glucose deprived (**OGD**)-BECs.

### 2.10. Measurement of mitochondrial function using Seahorse extracellular flux analysis

The mitochondrial function of recipient BECs treated with EVs during OGD conditions were evaluated using the Seahorse analysis by measuring oxygen consumption (**OCR**) (38). BECs seeded at 20,000 cells/well were cultured in a Seahorse XF96 plate for three days. Post-incubation, the cells were washed with PBS pH 7.4 and treated with OGD medium for four hours in a hypoxic chamber. The cells were incubated with SC-lEVs, DGC-lEVs, and NVEP-lEVs at different EV concentrations in OGD medium for 24 h. Post-incubation, the medium was replaced with pre-warmed DMEM supplemented with 35 mM glucose, 5 mM glutamine and 8 mM pyruvate and used for Seahorse analysis. After measurement of baseline OCR, 2.5 μmol/L oligomycin A and 0.7 μmol/L carbonyl cyanide-p-trifluoromethoxyphenyl-hydrazone were consecutively added to measure the proton leak and maximal OCR, respectively (17, 38).

### 2.11. Statistical analysis

Statistically significant differences between the means of controls and treatment groups or within treatment groups were determined using Student’s t-test, one-way, or two-way analysis of variance (**ANOVA**) with post hoc corrections using multiple comparison tests at α=0.05 using GraphPad Prism 10 (GraphPad Software). The notations for the different significance levels are indicated as follows: *p < 0.05, **p < 0.01, ***p < 0.001, ****p < 0.0001.

## 3. Results

### 3.1. Physicochemical characterization of lEVs isolated via SC and DGC

lEVs were isolated from conditioned medium of BECs by SC using our previously reported protocols (35, 36, 39–41). The SC-derived lEV fraction was further purified by high-resolution iodixanol DGC. **Figure 1** schematically illustrates the workflow for lEV isolation using SC and subsequent DGC purification. Briefly, EV-conditioned medium collected from confluent BECs were centrifuged at 300 ×g for 10 min to remove cell debris, followed by 2000 ×g for 20 min to pellet apoptotic bodies. The resulting supernatant was centrifuged at 20,000 ×g for 45 min to collect lEVs (**Fig. 1**). The resulting lEV pellet was resuspended in PBS and referred to as **SC-lEVs**. For DGC, SC-lEVs were mixed with iodixanol to achieve a final concentration of 36% and overlaid with step gradients of 30%, 24%, 18%, and 12% (w/v). The 12–36% (w/v) gradient was centrifuged at 120,000 ×g for 15 h (**Fig. 1**). The first seven fractions were pooled, diluted with PBS, and centrifuged at 120,000 ×g for 4 h. The resulting lEV pellet was resuspended in PBS and referred to as **DGC-lEVs**. The lower five fractions were designated as non-vesicular extracellular particles (**NVEPs**) and are referred to as NVEP-lEVs.

**Figure 1:**
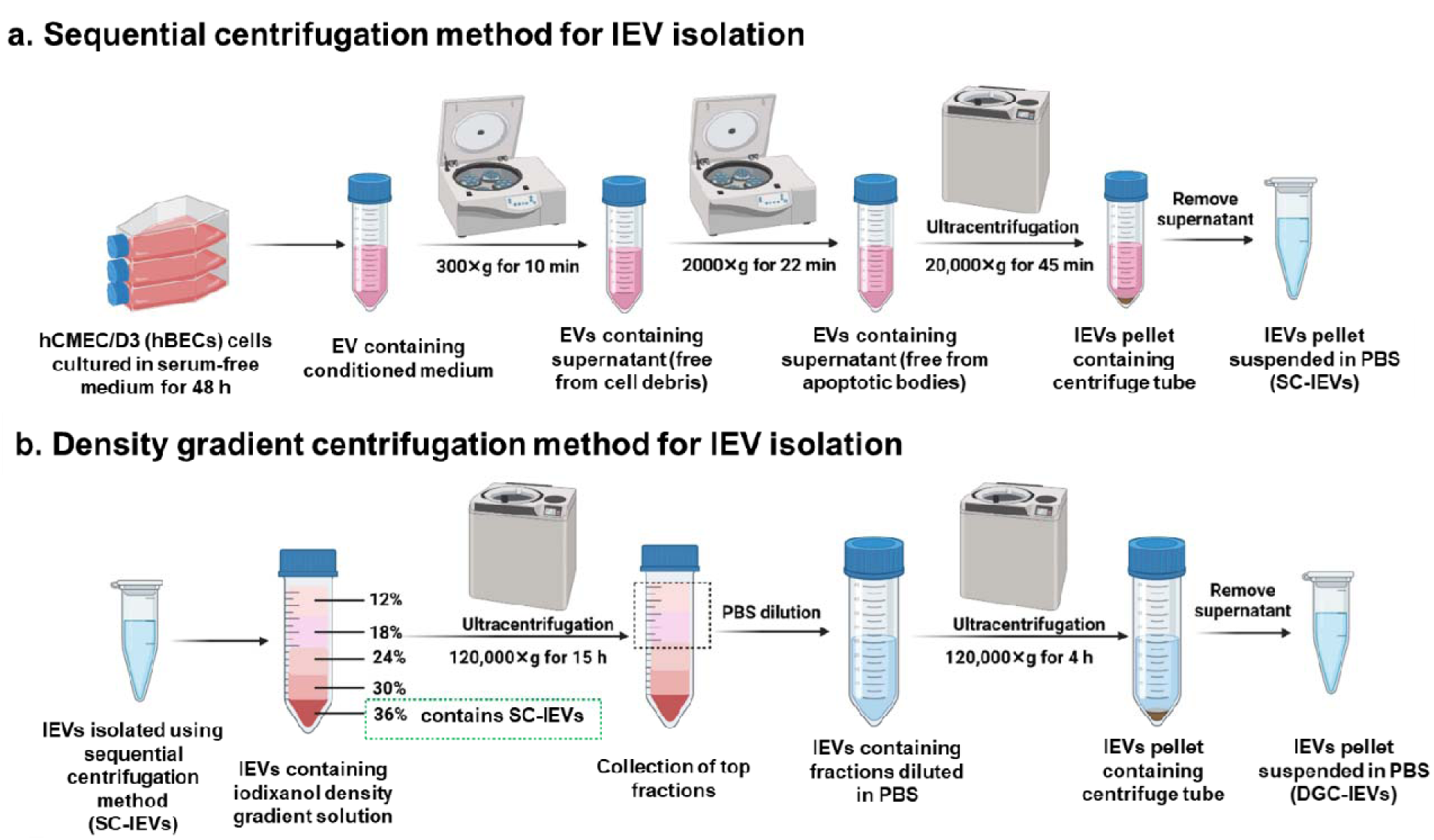
Schematic illustration of lEV isolation from conditioned medium of human brain endothelial cells. lEVs were first collected via sequential ultracentrifugation (**SC**) and subsequently purified using iodixanol density gradient centrifugation (**DGC**).

We first quantified and compared the protein yield of SC- and DGC-lEVs. Approximately 1,500 μg of SC-lEV protein yielded only 50–400 μg of DGC-lEV protein, corresponding to an average protein loss of ∼80–90% during DGC purification (**Fig. 2a**). This substantial decrease highlights the high stringency of the DGC process. In comparison, the retained NVEP fraction yielded ∼193 μg protein (∼7.7% recovery, **Supplemental Fig. 1**), indicating that a substantial proportion of the discarded material consisted of non-vesicular components rather than vesicle-enriched particles. Dynamic light scattering further revealed marked differences in physicochemical properties between SC- and DGC-lEVs. SC-lEVs exhibited an average diameter of ∼236 nm, consistent with previous reports (12–15, 42), whereas DGC-lEVs were significantly larger (∼512 nm, *p*<0.001; **Fig. 2b**). The particle size distribution shifted from 30–600 nm (peak ∼200 nm) for SC-lEVs to 200–800 nm (peak ∼400 nm) for DGC-lEVs, indicating selective depletion of smaller vesicles and non-vesicular particles during DGC purification (**Fig. 2c**). The dispersity index also increased from 0.4 (SC) to 0.5 (DGC; **Fig. 2d**). Importantly, dynamic light scattering analysis of the retained NVEP fraction revealed an average particle size of approximately 4 nm (**Supplemental Fig. 1b**). This size is several orders of magnitude smaller than extracellular vesicles and substantially below the dimensions expected for free mitochondria. These findings strongly suggest that the lower-density fractions do not contain a substantial population of large EVs or free mitochondria but instead are enriched in non-vesicular extracellular nanoparticles, such as exomeres and supermeres (26). In contrast, SC-lEVs, DGC-lEVs, and NVEPs exhibited average zeta potentials of approximately −24.8 mV,−20.7 mV, and −25.7 mV, respectively (**Fig. 2e and Supplemental Fig. 1c**), suggesting that iodixanol exposure and prolonged centrifugation do not alter lEV surface charge. Representative TEM images of negatively stained SC-lEVs showed heterogeneous vesicles ranging from 150–800 nm (**Fig. 2f**). The observed morphologies are consistent with TEM images of EVs in the published literature (43, 44).

**Figure 2:**
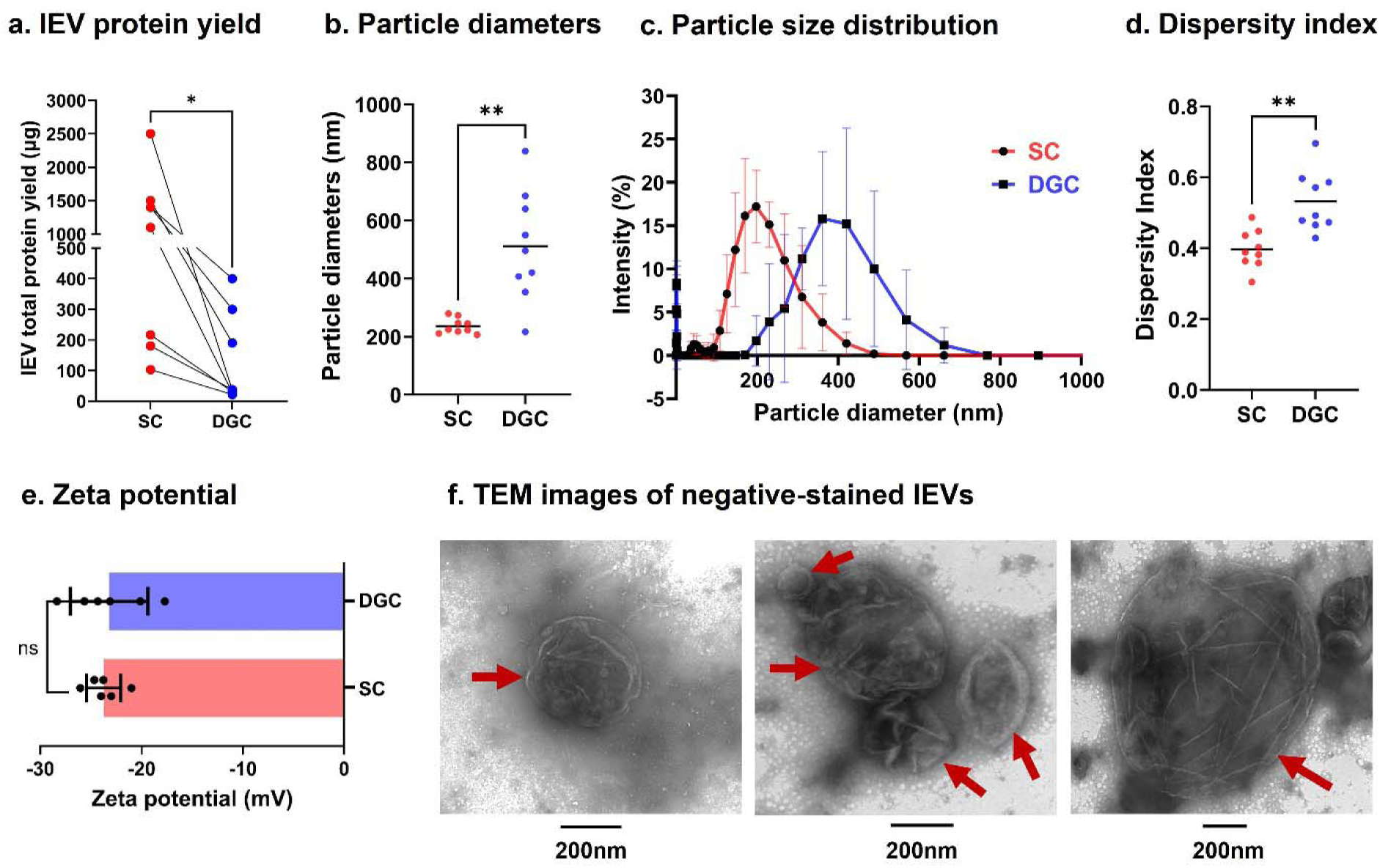
Physicochemical characterization of lEVs isolated by sequential centrifugation (SC) and purified by density gradient centrifugation (DGC). (**a**) Total lEV protein yield quantified by micro-BCA assay. (**b–e**) Dynamic light scattering (**DLS**) analysis of SC- and DGC-lEVs. Samples were diluted to 0.1 mg lEV protein/mL in PBS (pH 7.4) for particle size and dispersity index measurements and in deionized water for zeta potential measurements. DLS was performed using a Malvern Zetasizer Pro. (**f**) Representative negative-stain TEM images of SC-lEVs showing vesicle morphology (scale bar = 200 nm).

### 3.2. Detection of mitochondria and mitochondrial proteins in SC-lEVs

We acquired TEM images of cross-sectioned SC-lEVs to assess the presence of mitochondria within the vesicle lumen (**Fig. 3a**). The blue arrowhead denotes the lEV membrane, while maroon arrowheads highlight mitochondria. These appeared as membrane-bound, electron-dense structures within the lEV lumen. lEVs varied in size and carried differing numbers of mitochondria, which themselves exhibited diverse morphologies and dimensions. Notably, the mitochondrial structures observed here closely resemble extracellular free mitochondria and mitochondria-containing EVs described in previous reports (45–47). We performed a semi-quantitative analysis of TEM images obtained from cross-sectioned SC-lEVs. For the TEM analysis, uncropped TEM images were evaluated to identify total lEVs and mitochondria-containing lEVs within each imaging field (**Supplemental Fig. 2** and **Supplemental Table 2**). Mitochondria were identified as membrane-bound electron-dense structures enclosed within the vesicle lumen. Analysis of four independent TEM fields revealed that approximately 13.8% (8/58, **SL Fig. 2a**), 9.1% (6/66, **SL Fig. 2b**), 12.5% (1/8, **SL Fig. 2c**), and 8.0% (2/25, **SL Fig. 2d**) of SC-lEVs contained detectable mitochondria, corresponding to an average of 10.9% mitochondria-containing lEVs across all analyzed fields. Consequently, approximately 89.1% of SC-lEVs did not contain morphologically identifiable mitochondria (**Supplemental Table 2**). We further quantified the number of mitochondria per vesicle among the mitochondria-containing lEV population. Of these vesicles, 70.6% contained a single mitochondrion, 23.5% contained two mitochondria, and 5.9% contained three mitochondria (**Supplemental Table 2**), indicating that the majority of mitochondria-containing lEVs carry a single mitochondrion.

**Figure 3:**
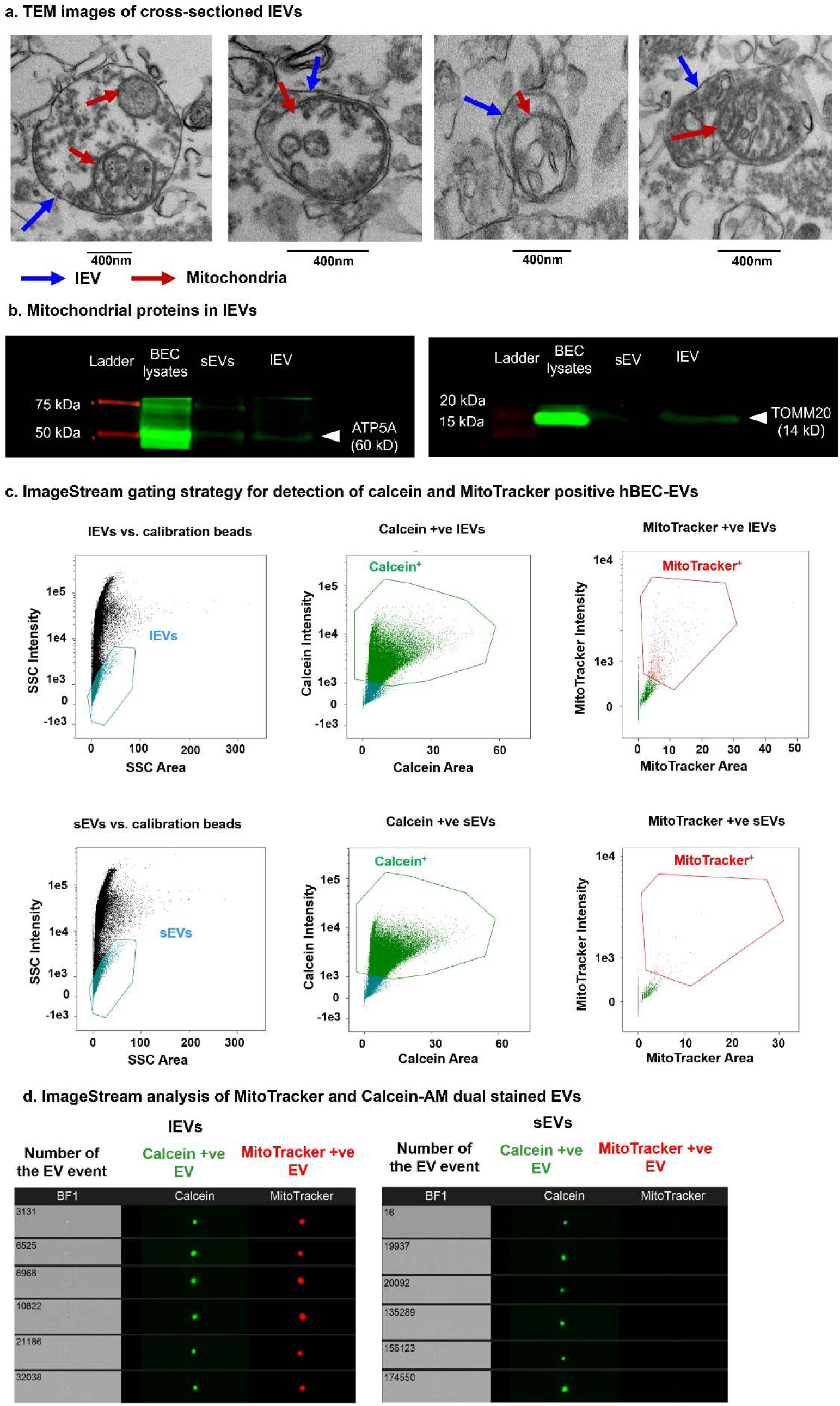
**Presence of mitochondria and mitochondrial proteins in lEVs isolated from SC. a) TEM imaged of cross-sectioned lEVs**: Representative TEM images of sectioned lEVs isolated from SC. lEVs (blue arrow) contained one or two mitochondria (electron-dense structures, maroon arrows) in the lEV lumen. Scale bar 400 nm. **b) Detection of mitochondrial proteins in lEVs using western blot analysis:** Western blots show the relative expression of ATP5A (54 kDa) and TOMM20 (14 kDa) in lEVs. Cell lysates and small EVs (sEVs) were used as a control, and protein standards were used as a reference for the molecular mass of protein bands. **c) Representative ImageStream gating strategy for detection of calcein and Mitotracker positive lEVs.** lEVs and sEVs were gated from calibration beads on SSC area *vs*. intensity plots. Total EV populations were gated for calcein-positive events in the green fluorescence channel. Total calcein-positive EVs were gated for MitoTracker-positive events in the red fluorescence channel. **d) ImageStream analysis of MitoTracker and Calcein-AM dual stained lEVs.** Calcein-AM and MitoTracker dual stained lEVs and sEVs were run through ImageStream flow cytometer. About 100,000 total events were captured in the SSC area versus SSC intensity plots where calcein-positive EVs (green fluorescence, channels Ch02) were gated to separate all events from the calibration beads. A total of 5,000 calcein-positive events were acquired for further gating MitoTracker-positive events (red fluorescence) on red fluorescence channels. Single color stains (i.e., only calcein+ve EVs and only MitoTracker+ve EVs) and calibration beads were used to adjust spectral compensations.

We evaluated the presence of structural and functional mitochondrial proteins in SC-lEVs by western blotting. As a control, we used small EVs (**sEVs**/exosomes) were isolated at 120,000 ×g for 90 min using SC. Both sEVs and lEVs were analyzed for ATP5A (a 60 kDa subunit of ATP synthase in the electron transport chain) and TOMM20 (a 14 kDa subunit of the translocase of the outer mitochondrial membrane complex). ATP5A was detected in both sEVs and lEVs, whereas TOMM20 was selectively present in lEVs, with no detectable 14 kDa band in sEVs (**Fig. 3b**). These findings indicate that lEVs harbor functional (ATP5A) and structural (TOMM20) mitochondrial components, consistent with mitochondria within SC-lEVs confirmed using TEM (**Fig. 3a**).

Orthogonally, the presence of mitochondria in SC-lEVs was confirmed using ImageStream analysis, which integrates flow cytometry with fluorescence imaging to enable direct visualization of mitochondria-positive lEVs (**Fig. 3c–d**). We demonstrated sEVs as a negative control to demonstrate the selective presence of mitochondria in lEVs. SC-sEVs and lEVs were isolated from BECs pre-stained with MitoTracker Deep Red, and then labeled with calcein-AM. Upon incubation, calcein-AM permeates the EVs and is hydrolyzed by intraluminal esterases to generate the membrane-impermeant green, fluorescent calcein (34, 48). Dual-stained EVs were analyzed by ImageStream to acquire calcein /MitoTracker EV images. The gating strategy (**Fig. 3c**) involved: (i) gating EVs from calibration beads on SSC area *vs*. intensity plots, (ii) selecting calcein EVs in the green fluorescence channel, and (iii) further gating for MitoTracker events in the red fluorescence channel. Representative images (**Fig. 3d**) demonstrate that calcein BEC-lEVs also exhibit MitoTracker signals, indicating the presence of mitochondrial components. These findings are consistent with published reports showing MitoTracker-positive signals in CD63-positive EVs isolated from bronchial alveolar lavage fluid (37).

ImageStream-based quantification of Calcein-AM-positive EVs and Calcein-AM/MitoTracker double-positive EVs provided quantitative assessment of mitochondrial content within EV subpopulations. For SC-lEVs, a total of 81,657 events were analyzed, of which 53,971 events (66.1%) were Calcein-positive, confirming their vesicular identity (**Supplemental Table 1**). Among these EVs, 446 events were positive for both Calcein and MitoTracker, indicating the presence of a distinct mitochondria-containing EV subpopulation. In contrast, analysis of 132,937 SC-sEV events identified 93,774 Calcein-positive events (70.5% EVs), but only 37 Calcein/MitoTracker double-positive events. Importantly, SC-lEVs exhibited a ∼20.8-fold higher frequency of mitochondria-containing EVs compared with SC-sEVs. These quantitative findings are consistent with our qualitative ImageStream observations and provide additional evidence that mitochondria are predominantly associated with the lEV population rather than the sEV population.

### 3.3. DGC-lEVs outperform SC-lEVs in increasing recipient ischemic BEC bioenergetics

Mitochondria are central to cellular bioenergetics, synthesizing ATP from ADP via aerobic respiration. To evaluate the effects of mitochondria-containing EVs, we measured relative ATP levels in recipient BECs treated with sEVs, lEVs, and NVEPs using a luciferase-based ATP assay. Recipient BECs were exposed to oxygen-glucose deprivation (**OGD**) for 4 h in a hypoxic chamber, followed by treatment with lEVs isolated via SC or DGC at varying concentrations for 24 h. ATP levels in cells maintained in complete growth medium under normoxic conditions served as a control, while untreated OGD-exposed BECs were used as the OGD control (**Fig. 4a**).

**Figure 4:**
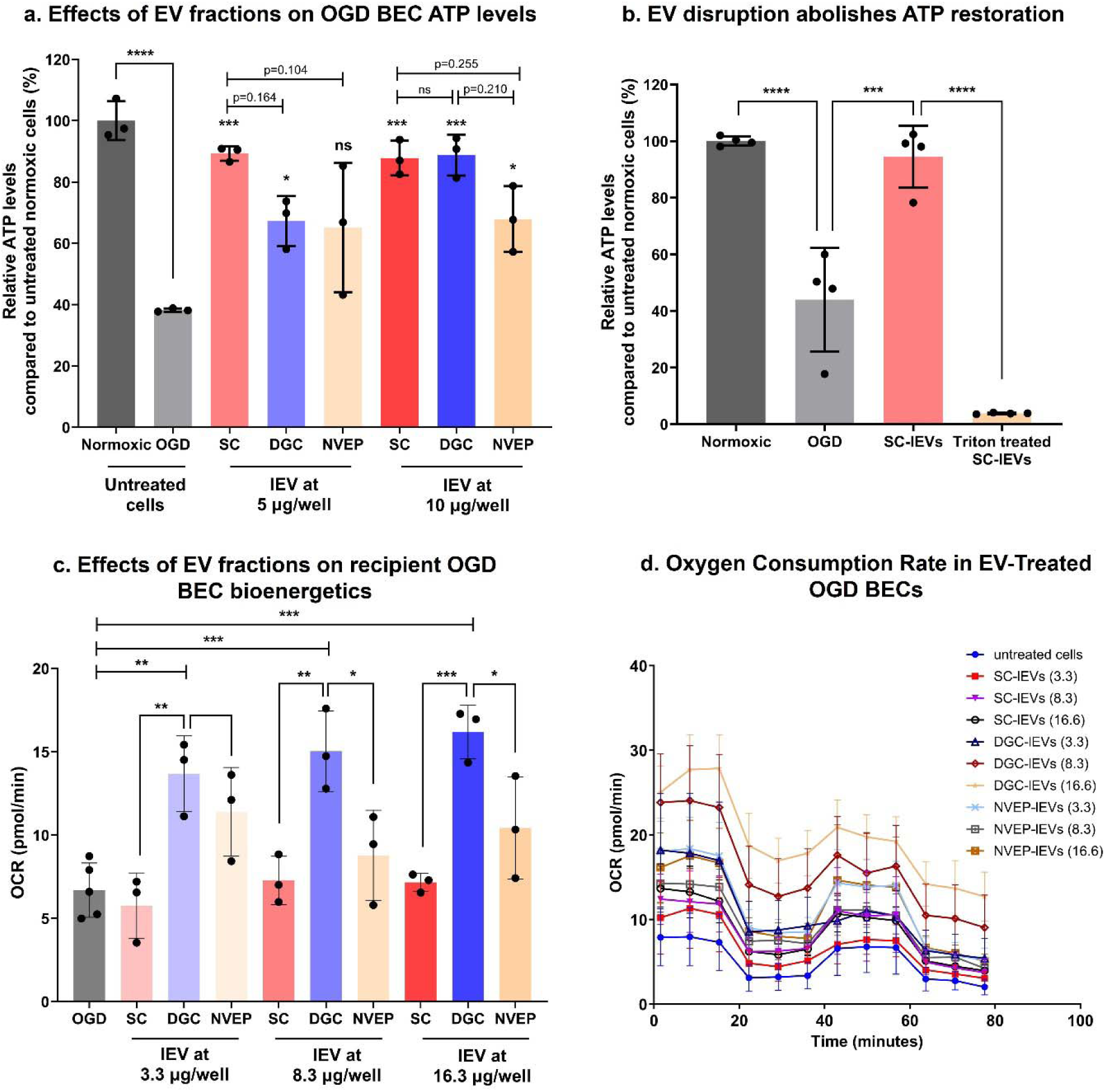
Effect of SC and DGC-derived lEVs on relative ATP levels and cellular bioenergetics in the recipient ischemic BECs. *(a) Effect of SC- and DGC-derived lEVs on relative ATP levels in the recipient ischemic BECs.* Recipient BECs were treated with BEC-derived lEVs isolated from SC or DGC or NVEP fractions at 5, and 10 µg EV protein/well for 24 h in OGD medium. Untreated cells incubated in the OGD medium were used as OGD control, whereas untreated cells incubated in complete growth medium were used as the normoxic control. Post-incubation, CellTiter-Glo 2.0 reagent was added to an equal volume of cell culture medium in each well. Luminescence was measured at 1 s integration time using a SYNERGY HTX multi-mode plate reader. The % relative ATP levels of the treated cells were calculated as follows = (relative luminescence unit (RLU) of treated cells/RLU of untreated OGD cells) × 100. The differences in ATP levels of the treated groups were compared against control using one-way ANOVA using GraphPad Prism software. *(b) Intact EV and mitochondria membranes required for their functional effects in increasing cellular ATP levels.* Recipient BECs were treated with BEC-derived SC-lEVs and Triton-X-100 treated SC-lEVs at 10 µg EV protein/well for 24 h in OGD medium. Untreated cells incubated in the OGD medium were used as OGD control, whereas untreated cells incubated in complete growth medium were used as the normoxic control. *(c-d) Effect of SC-lEVs and DGC-lEVs on recipient BEC mitochondrial respiration under OGD conditions.* BEC were cultured in a Seahorse XF96 plate for four days at 20,000 cells/well. Confluent BECs were incubated with OGD medium in a hypoxic chamber for four hours. Post-OGD exposure, cells were incubated with SC-lEVs, DGC-lEVs, and NVEP-lEVs at indicated doses diluted in OGD medium for 24 h. Post-treatment, the medium was replaced with DMEM supplemented with 35 mM glucose, 5 mM glutamine and 8 mM pyruvate. Oxygen consumption rate (**OCR**, **c**) and OCR profile (**d**) in BEC was analyzed using the Seahorse XFe96 analyzer. Data represents mean±SD (n=3). *p<0.05, ***p<0.001, ****p < 0.0001, ns: non-significant.

OGD exposure resulted in a marked reduction in ATP levels (∼60% decrease relative to normoxic controls; p < 0.0001), confirming substantial bioenergetic impairment (**Fig. 4a**). Treatment with low-dose SC-lEVs (5 μg/well) or DGC-lEVs significantly increased ATP levels compared with OGD controls, demonstrating that both preparations retained biological activity. Notably, SC-lEVs produced a greater ATP recovery than DGC-lEVs at this lower dose (**Fig. 4a**). SC-lEVs-mediated restoring cellular bioenergetics, consistent with our prior reports demonstrating functional mitochondrial delivery via SC-isolated EVs (12–15, 42). In contrast, NVEPs did not significantly improve ATP levels at the same concentration (**Fig. 4a**). At higher doses, both SC-lEVs and DGC-lEVs robustly restored cellular ATP levels, producing approximately a 2.3-fold increase relative to OGD controls (p < 0.0001), with no significant difference observed between the two groups (**Fig. 4a**). Although NVEP treatment at 10 μg/well also resulted in a significant increase in ATP levels, the magnitude of recovery remained approximately 20% lower than that achieved by either SC-lEVs or DGC-lEVs. These findings indicate that the vesicle-enriched fractions possess superior bioenergetic activity compared with the non-vesicular fraction.

To further investigate whether the bioenergetic effects of SC-lEVs depend on intact vesicular structures, we performed membrane disruption studies using Triton X-100. SC-lEVs were treated with 0.1% Triton X-100 for 30 min to disrupt membrane integrity, followed by dilution in PBS and centrifugation at 20,000 ×g for 45 min. The resulting pellet was resuspended in ice-cold PBS and used for subsequent functional studies. Consistent with our previous findings, treatment of OGD-exposed BECs with intact SC-lEVs significantly increased cellular ATP levels compared with untreated OGD controls. In contrast, Triton X-100-treated SC-lEVs completely lost their ability to restore cellular ATP levels, resulting in ATP values comparable to untreated OGD cells. These findings demonstrate that membrane integrity is essential for the bioenergetic activity of SC-lEVs and indicate that disruption of membrane structures abolishes their functional effects in recipient cells.

To further evaluate the functional effects of SC-lEVs, DGC-lEVs, and NVEPs on recipient cell bioenergetics using an orthogonal approach, we measured cellular oxygen consumption rates (**OCR**) using Seahorse extracellular flux analysis, a highly sensitive and widely accepted method for assessing mitochondrial oxidative phosphorylation [1-3]. OGD-exposed BECs were treated with SC-lEVs, DGC-lEVs, or NVEPs at 3.3, 8.3, and 16.5 μg/well for 24 h prior to OCR measurements. The selected doses correspond to 10, 25 and 50 μg EV protein/0.32 cm^2^ growth area per well in a standard 96-well plate. The growth surface area per well in a Seahorse plate is 0.106 cm^2^/well and we scaled-down the doses accordingly. Interestingly, treatment with SC-lEVs did not significantly increase OCR at any tested dose compared with untreated OGD controls. In contrast, DGC-lEV treatment resulted in a dose-dependent and significant increase in OCR, indicating enhanced mitochondrial respiratory activity in recipient BECs. Notably, NVEP treatment also failed to significantly increase OCR relative to OGD controls. Collectively, these findings demonstrate that DGC-lEVs outperform both SC-lEVs and NVEPs in promoting mitochondrial oxidative phosphorylation and respiratory function in recipient BECs.

To investigate whether the SC-lEV-mediated increase in cellular ATP was due to the transfer of innate lEV mitochondria, we labeled SC-lEV mitochondria with MitoTracker Red (MitoT-red-lEVs) prior to isolation from donor BECs. MitoT-red selectively stains polarized, functional mitochondria, allowing direct visualization of mitochondrial transfer. Recipient BECs were incubated with MitoT-red-lEVs at varying doses, and intracellular MitoT-red signals were assessed 48 h post-exposure using fluorescence microscopy. Untreated BECs exhibited no signal in the Cy5 channel, confirming assay specificity. Recipient BECs treated with 10 μg MitoT-red-lEVs displayed clear intracellular mitochondrial signals, which further intensified at higher doses (30 and 60 μg EV protein/well; **Fig. 5**). These findings provide a mechanistic basis for the ATP assay results, showing that SC-lEVs transfer mitochondria into recipient BECs, thereby directly contributing to the observed dose-dependent enhancement of relative cellular ATP levels under OGD conditions.

**Figure 5:**
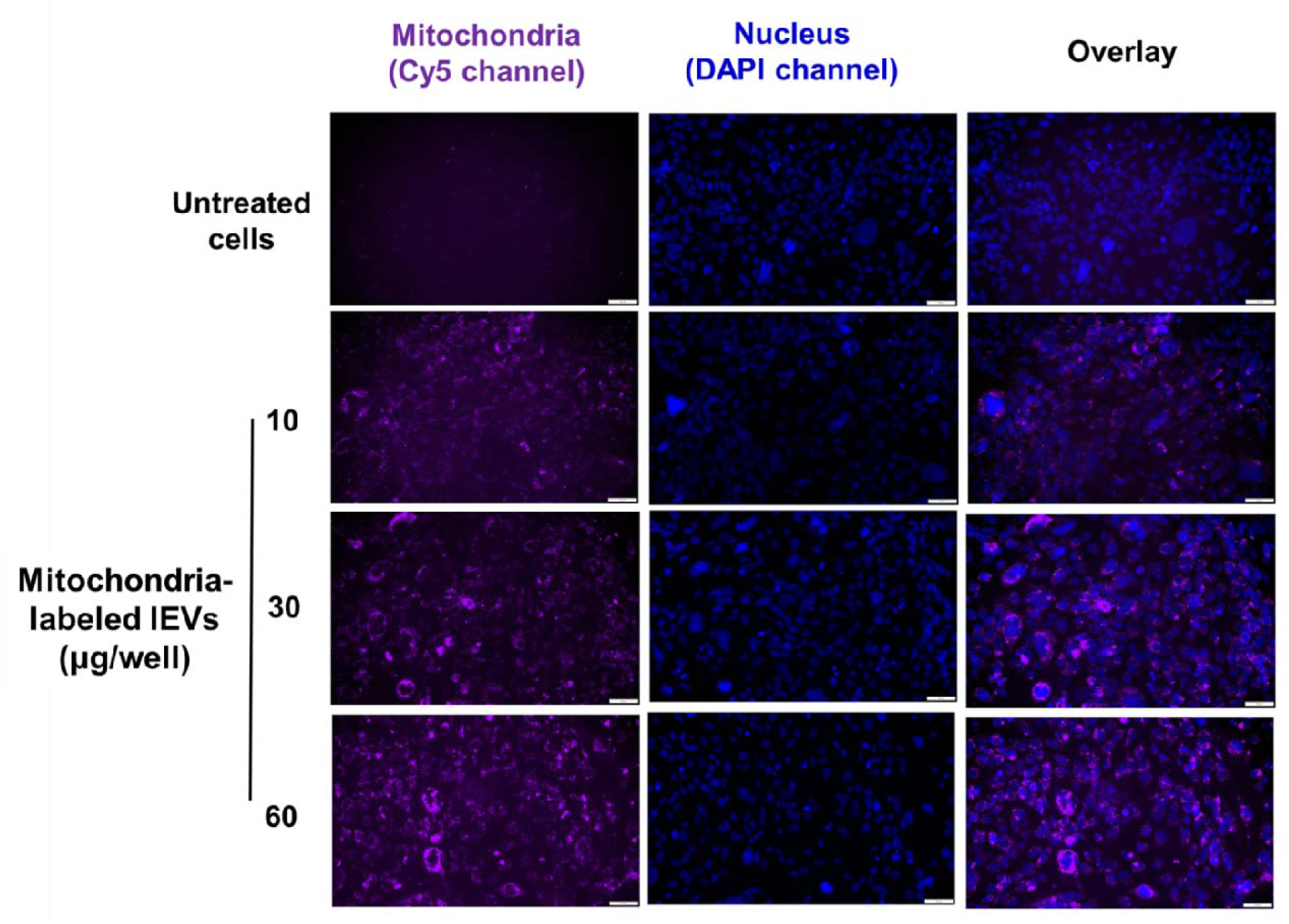
lEV mitochondria transfer into recipient BECs at varying doses. BECs were cultured in 48-well plates in a humidified incubator at 37 °C. Recipient BECs were incubated with indicated amounts of MitoT-red-lEV diluted in complete growth medium for 48 h. Cells treated with complete growth medium were used as control, untreated cells. Post-incubation, the cells were washed and incubated with the phenol-red medium. MitoT-red signals (purple puncta) and nucleus (blue) were observed under an Olympus IX 73 epifluorescent inverted microscope using Cy5 (pseudo purple colored for mitochondria) and DAPI (pseudo blue colored for nucleus) channels at 20× magnification—scale bar 50 μm.

## 4. Discussion

Ischemic stroke–induced OGD damages mitochondria in BECs, leading to brain injury, BBB disruption, and long-term cognitive deficits (49). Protecting mitochondrial function via exogenous mitochondrial supplementation is, therefore, a promising strategy to enhance BEC survival after stroke. In our prior comprehensive investigations, we extensively characterized both sEVs and lEVs using multiple orthogonal approaches (35, 36, 39, 40). These studie demonstrated that lEVs contain mitochondria and mitochondrial proteins, whereas sEVs lack detectable mitochondria, consistent with previously published reports. Importantly, lEVs, but not sEVs, transferred mitochondria to recipient brain endothelial cells (**BECs**), resulting in increased relative cellular ATP levels and restoration of bioenergetics in oxygen-glucose deprived (**OGD**) BECs. Furthermore, intravenous administration of mitochondria-containing lEVs significantly reduced infarct volume and improved neurological outcomes in a mouse model of ischemic stroke, whereas sEVs did not exhibit comparable mitochondria-mediated therapeutic effects (36, 39). Given these prior findings, the current study was specifically designed to further optimize and characterize mitochondria-enriched lEV preparations. In our previous works, we have isolated lEVs by SC, which likely yields a heterogeneous mixture of vesicles with and without mitochondria. Because the therapeutic effect largely depends on mitochondria-containing lEVs, their separation from other vesicles is critical for developing effective lEV mitochondria–based therapies. This study examined whether purifying SC-lEVs can enhance their therapeutic benefits in ischemic BECs. We compared SC-lEVs and DGC-purified SC-lEVs for their physicochemical properties and functional efficiency. SC-lEVs contained a heterogeneous mixture of vesicles (30–600 nm), whereas DGC enriched for larger vesicles (200–600 nm) while removing smaller particles and contaminants—presumably enriching mitochondria-packed lEVs but at the cost of substantial yield loss. Functionally, SC-lEVs and DGC-lEVs both significantly restored ATP levels in OGD-injured BECs with no difference between groups. However, OCR revealed distinct functional effects: SC-lEVs did not significantly alter respiration, whereas DGC-lEVs induced a dose-dependent increase in OCR, indicating enhanced oxidative phosphorylation. These findings further support that DGC purification yields a more mitochondria-enriched and functionally potent lEV preparation with enhanced capacity to support oxidative phosphorylation in recipient cells.

DGC separates lEVs based on buoyant density using iodixanol gradients (12–36%). We determined the density of each fraction collected from the iodixanol gradient (**Supplemental Table 3**). The average densities of the collected fractions ranged from 1.027 g/mL in fraction 1 to 1.217 g/mL in Fraction 10. Importantly, the top seven fractions retained as DGC-lEVs in the revised study exhibited densities ranging from 1.027 to 1.102 g/mL (Fractions 1–7). These density values are consistent with the broad density range previously reported for large extracellular vesicles and microvesicle-like populations isolated by iodixanol gradient centrifugation. lEVs typically have a buoyant density ranging between 1.10–1.19 g/mL (22), with mitochondria-containing lEVs occupying the lower end of this range. During centrifugation, the gradient stratifies particles: the upper layers (12–20% iodixanol, corresponding to densities of approximately 1.06–1.11 g/mL, (32, 50)) align with the density of mitochondria-containing lEVs. This allows these vesicles to migrate and concentrate in these less dense layers, separating them from other components. The mid-layer gradients, around 24% iodixanol (∼1.12–1.15 g/mL, (32, 50)), and likely capture standard lEVs without mitochondria. Meanwhile, the denser lower layers (30–36% iodixanol, ∼1.16–1.22 g/mL, (32, 50)) trap smaller, denser particles such as supermeres, lipoproteins, and other non-vesicular contaminants (26). Interestingly, the lower fractions in our DGC preparations showed average particle diameters of around <200 nm and an average zeta potential of -26 mV suggestive of non-vesicular contaminants.

Notably, intact/free mitochondria have been reported to exhibit substantially higher buoyant densities, generally exceeding ∼1.16 g/mL and frequently sedimenting within higher-density regions of iodixanol gradients. This value corresponds to intact, full-length mitochondria isolated by DGC; however, full-length mitochondria are unlikely to be incorporated into extracellular vesicles due to size and biogenesis constraints, and EV-associated mitochondrial material is more likely to consist of fragments or partially functional mitochondrial components rather than whole organelles (51). Consistent with these observations, the highest-density fractions in our gradients (Fractions 8–10; 1.170–1.217 g/mL) were not included in the DGC-lEV preparation. Importantly, these fractions were retained in the revised study and independently characterized as NVEPs. We subsequently characterized the lower-gradient material retained during the revised study and found that it was compositionally and functionally distinct from the vesicle-enriched DGC-lEV fraction. We would also like to clarify that our rationale for collecting the upper seven fractions was to enrich EVs, including mitochondria-containing lEVs, while minimizing the incorporation of higher-density non-vesicular material. The objective of the DGC procedure was not to maximize recovery of total mitochondrial material, but rather to enrich vesicle-associated mitochondrial cargo. Extending fraction collection into progressively denser regions of the gradient would likely increase co-isolation of non-vesicular extracellular particles, protein aggregates, free organellar material, and other dense contaminants, thereby reducing the specificity and purification advantage afforded by density-gradient centrifugation.

We compared lEV yield and particle size between SC- and DGC-EVs. DGC retained only 17% of the total protein from the initial SC sample, indicating substantial loss that complicates downstream characterization and functional studies. This pronounced reduction aligns with prior reports: Brennan *et al.* showed that small EVs from 200 μL human serum dropped from 772 μg (SC) to 31.1 μg (DGC), with particle concentrations falling from 6.3 × 10 to 4 × 10 particles/mL (22). Similarly, Dhondt *et al.* reported only 30% EV recovery from prostate cancer patient urine via DGC compared to ultracentrifugation (52).

SC-isolated lEVs averaged ∼250 nm with a broad 30–600 nm size distribution (**Fig. 2b, c**). Our lEV preparations exhibited polydispersity indices ranging from 0.4–0.5 (**Fig. 2d**), indicating a broad size distribution rather than a uniform vesicle population. DGC purification increased the mean size to 512 nm by removing smaller vesicles and non-vesicular particles, selectively enriching lEVs in the 200–600 nm range (**Fig. 2c**). Specifically, negative-stain TEM revealed lEVs ranging from approximately 200–800 nm in diameter. In addition, TEM analysis of sectioned EVs showed that mitochondria-containing lEVs were predominantly present in vesicles larger than 300 nm, with diameters ranging from approximately 400–800 nm. The observed differences primarily reflect the heterogeneous nature of the lEV population as well as the distinct measurement principles of these techniques. Importantly, the DLS measurements represent the average hydrodynamic diameter of the entire vesicle population in suspension, whereas TEM directly visualizes individual vesicles. Consequently, DLS reflects the collective contribution of both smaller and larger vesicles (that is important to account for), while TEM reports the size of specific vesicles captured within the imaging field. This distinction is particularly relevant for heterogeneous EV samples, where a relatively small subpopulation of large mitochondria-containing lEVs coexists with a larger population of smaller vesicles. Consistent with this interpretation, the DGC-lEV preparation exhibited a z-average diameter of approximately 500 nm, and its particle size distribution indicated that ∼75% of vesicles were larger than 300 nm, supporting the enrichment of larger mitochondria-containing lEVs following DGC purification. Furthermore, DLS and TEM inherently operate on different measurement principles. DLS estimates the hydrodynamic diameter of particles in their hydrated state based on Brownian motion and light scattering, whereas TEM measures the physical dimensions of individual vesicles after sample preparation and imaging under vacuum. Sample processing, dehydration effects, and the limited number of vesicles visualized by TEM can all contribute to differences in measured particle sizes. Therefore, direct numerical agreement between DLS and TEM measurements is not necessarily expected, particularly for highly heterogeneous EV populations.

Brennen *et al.* demonstrated that DGC is not only effective in improving the purity of EVs but also influences the size distribution of isolated particles compared to SC (22). The study reported a significant increase in the average particle diameter of EVs after DGC purification, with SC-isolated EVs having an average size of 93 nm (22). In contrast, DGC-purified EVs showed a larger average diameter of 131 nm (22). Size- and density-based separation is crucial for enriching mitochondria-containing lEVs, which TEM analysis showed consistently exceed 200 nm, with a minimum observed size of 400 nm (**Fig. 3a**). Larger lEVs often contained more or larger mitochondria, indicating that mitochondrial content drives particle size. By removing smaller vesicles and non-vesicular particles, DGC enhances the purity and functional relevance of these biologically distinct lEVs, providing a robust platform to study their bioenergetic and therapeutic potential. Before comparing functional benefits of SC- and DGC-lEVs, we confirmed that SC-lEVs contain mitochondria and mitochondrial proteins. TEM of cross-sectioned lEVs revealed electron-dense structures in the lumen (**Fig. 3a**), and western blots detected ATP5A and TOMM20 (**Fig. 3b**), consistent with prior reports (20, 45). Ikeda *et al.* showed the presence of ATP5A proteins in human cardiomyocyte-derived mitochondria-containing lEVs (45), whereas Silva *et al.* showed human mesenchymal stromal cell-derived lEVs expressed TOMM20 (20). Additionally, ImageStream analysis of calcein- and MitoTracker-labeled BEC-EVs further confirmed mitochondrial components in SC-lEVs but not SC-sEVs (**Fig. 3d**). We were unable to perform additional ImageStream analyses on DGC-lEV preparations due to restricted sample quantities and the need to use those samples for functional activity assays (**Fig. 4**). Therefore, a rigorous comparison of particle sizes among SC-lEVs and DGC-lEVs using ImageStream was not feasible. However, future study will involve systematic comparisons of DGC-purified lEVs *vs*. sEVs using ImageStream analysis.

Due to the substantial loss of lEVs during DGC purification, TEM, western blot, and ImageStream analyses were not feasible for DGC-lEVs. TEM imaging of cross-sectioned vesicles requires a high yield with a visible pellet; however, DGC resulted in ∼82–87% loss of SC-lEVs, making repeated attempts impractical. Similarly, western blot analysis typically requires 50–100 µg of protein to detect mitochondrial markers (ATP5A, TOMM20) and EV markers. DGC-lEVs consistently yielded only ∼25 µg of highly diluted protein, insufficient for reliable detection of these proteins. ImageStream analysis also necessitates concentrated EV preparations, which cannot be achieved with the low concentration of DGC-lEVs obtained. Despite these limitations, multiple lines of evidence strongly support the presence of mitochondria-containing EVs in DGC-lEVs. TEM analysis of SC-lEVs revealed mitochondria within lEVs >400 nm (**Fig. 3a**), and our DLS data show that DGC-lEVs predominantly comprise particles in the 400–700 nm range (**Fig. 2b and c**). This indicates that the larger, mitochondria-containing SC-lEVs are retained in the DGC fractions. Future studies aimed at isolating larger quantities of DGC-lEVs will enable detailed structural characterization by TEM, quantitative protein analysis by western blotting, and ImageStream analysis to further validate the presence and functional integrity of mitochondria in these vesicles.

Mitochondrial oxidative phosphorylation generates 80–90% of cellular ATP, which drops significantly under OGD. Four hours of OGD reduced BEC ATP levels by ∼60% compared to normoxic controls (**Fig. 5**), consistent with previous reports (53, 54). After OGD exposure, BECs were treated with SC- or DGC-lEVs at varying doses for 24 h. SC-lEVs and DGC-lEVs both significantly restored ATP levels in OGD-injured BECs (∼2.3-fold vs control, p < 0.0001) with no difference between groups (**Fig. 4a**). O’Brien *et al.* demonstrated that human MSC-derived lEVs transfer mitochondria to human cardiomyocytes, colocalizing with endogenous mitochondria, boosting ATP levels, and reducing apoptosis in doxorubicin-injured cells (44). Similarly, Silva *et al.* reported a three-fold increase in ATP levels and a six-fold increase in mitochondrial respiration in human pulmonary microvascular endothelial cells treated with MSC-derived lEVs (20). However, OCR measured using Seahorse analysis revealed distinct functional effects: SC-lEVs did not significantly alter respiration, whereas DGC-lEVs induced a dose-dependent increase in OCR, indicating enhanced oxidative phosphorylation. Importantly, these Seahorse findings should be interpreted together with the ATP assay results. Although both SC-lEVs and DGC-lEVs significantly restored ATP levels in OGD-injured BECs, the two preparations exhibited distinct effects on mitochondrial respiration. This observation is biologically plausible because ATP content and OCR represent related but fundamentally different measures of cellular bioenergetics. OCR specifically reflects mitochondrial oxidative phosphorylation, whereas relative cellular ATP levels were measured using a CellTiter-Glo luminescence assay, in which relative luminescence units (**RLU**) provide a signal proportional to ATP content in viable cells rather than an absolute quantification of intracellular ATP. Thus, RLU values reflects a luminescent readout that represent the overall energetic state of the cell and can be influenced by multiple mechanisms, including mitochondrial transfer, transfer of mitochondrial proteins, enhancement of glycolytic pathways, or other pro-survival signaling effects. One potential explanation for the increased OCR in DGC-lEV-treated cells is the compositional difference between SC-lEV and DGC-lEV preparations. SC-lEVs represent a heterogeneous population containing mitochondria-containing vesicles together with co-isolated extracellular proteins and other NVEPs. Consequently, when equal protein doses are administered, SC-lEV preparations likely contain fewer mitochondria-containing vesicles than DGC-lEV preparations. In contrast, DGC purification enriches for larger vesicles and mitochondria-containing lEVs while removing a substantial proportion of co-isolated material. Therefore, at equivalent protein doses, DGC-lEVs likely deliver a greater quantity of functional mitochondrial cargo to recipient cells, resulting in a more pronounced enhancement of oxidative phosphorylation.

This interpretation is further supported by the ATP and OCR data obtained from the NVEP fraction. Although NVEPs produced a modest increase in cellular ATP levels, they failed to enhance OCR. Given that NVEPs exhibited an average particle size of approximately 4 nm (**Supplemental Fig. 1b**) and lacked detectable vesicular or mitochondrial structures, these findings suggest that non-vesicular extracellular components may contribute to partial ATP recovery without directly improving mitochondrial respiratory function. Thus, ATP restoration alone does not necessarily indicate enhancement of oxidative phosphorylation. Taken together, the ATP and Seahorse datasets are complementary rather than contradictory. The ATP assay demonstrates that both SC-lEVs and DGC-lEVs improve the overall bioenergetic status of OGD-injured BECs, whereas Seahorse analysis provides mechanistic insight into mitochondrial respiratory activity. The combined findings indicate that DGC purification enriches for a functionally active mitochondria-containing lEV population with superior capacity to enhance oxidative phosphorylation, while SC-lEVs improve cellular energetics through a combination of mitochondrial and non-mitochondrial bioactive cargo. These results further support the conclusion that DGC purification yields a more mitochondria-enriched and functionally potent lEV preparation.

## 5. Conclusion

Delivering mitochondria-containing lEVs to ischemic BECs is a promising strategy to enhance cellular bioenergetics following stroke. Both SC- and DGC-derived lEVs restored ATP levels in injured BECs, while DGC-enriched lEVs demonstrated superior mitochondrial respiratory activity, indicating enrichment of a more functionally potent mitochondria-containing EV population. These findings identify DGC as a robust approach for isolating bioenergetically active lEVs and support the development of EV-based therapies for diseases associated with mitochondrial dysfunction.

## 6. Acknowledgements

The authors gratefully acknowledge National Institutes of Health grant number 7R01NS136752-02 for funding support. We also thank Mr. Nevil Abraham (Flow core, University of Pittsburgh) for assistance with ImageStream analysis.

There are no conflicts of interest to declare.

