## Supplemental File for "Sequential- *vs*. density gradient- centrifugation for the isolation of mitochondria-containing extracellular vesicles"

6431 Fannin Street, MSE R274

Houston, TX 77030

ORCID: 0000-0002-6363-3679

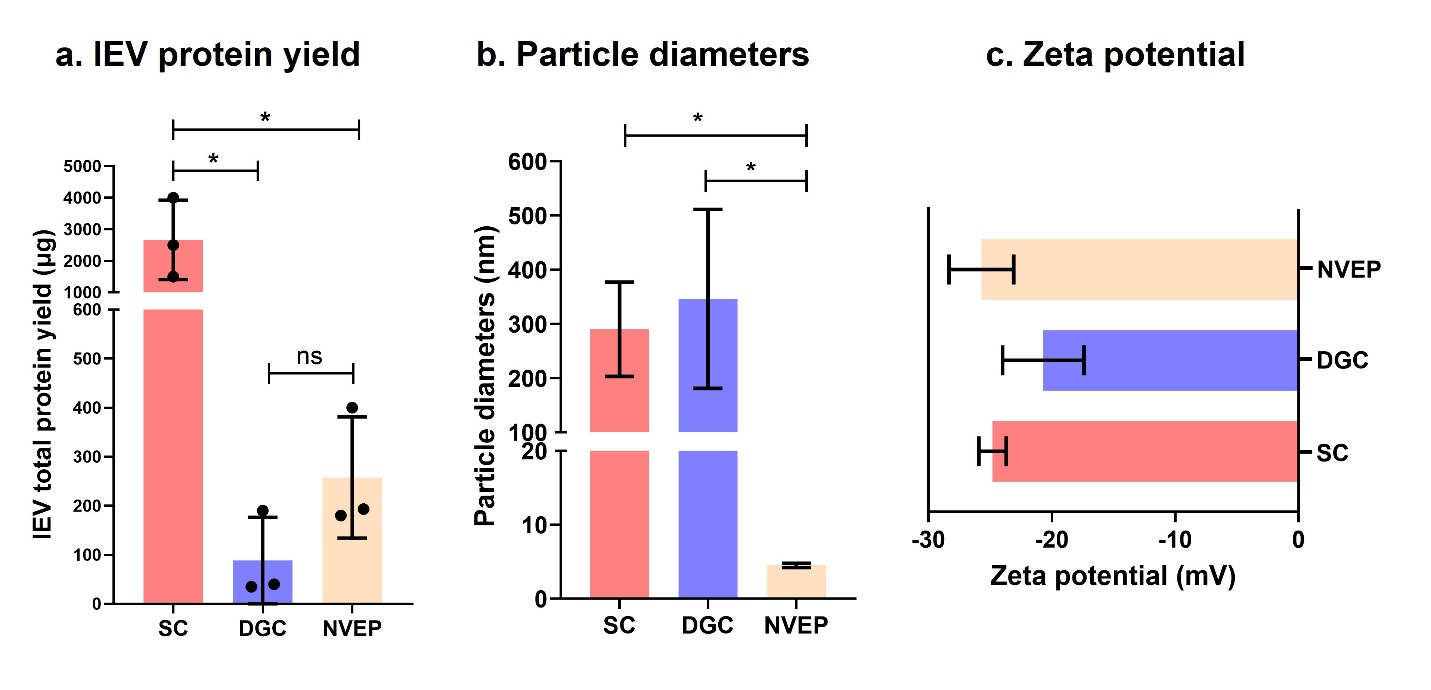

**Supplemental Figure 1: Physicochemical characteristics of lEVs isolated from SC- and DGC-methods as well as non-vesicular extracellular particles (NVEP) fractions obtained as a by-product of DGC-based purification. (a)** Total lEV protein yield quantified by microBCA assay. **(b-c)** Dynamic light scattering (**DLS**) analysis of SC- and DGC-lEVs. Samples were diluted to 0.1 mg lEV protein/mL in PBS (pH 7.4) for particle size measurements and in deionized water for zeta potential measurements. DLS was performed using a Malvern Zetasizer Pro. Data represents mean±SD (n=3). * p<0.05, ns= not significant.

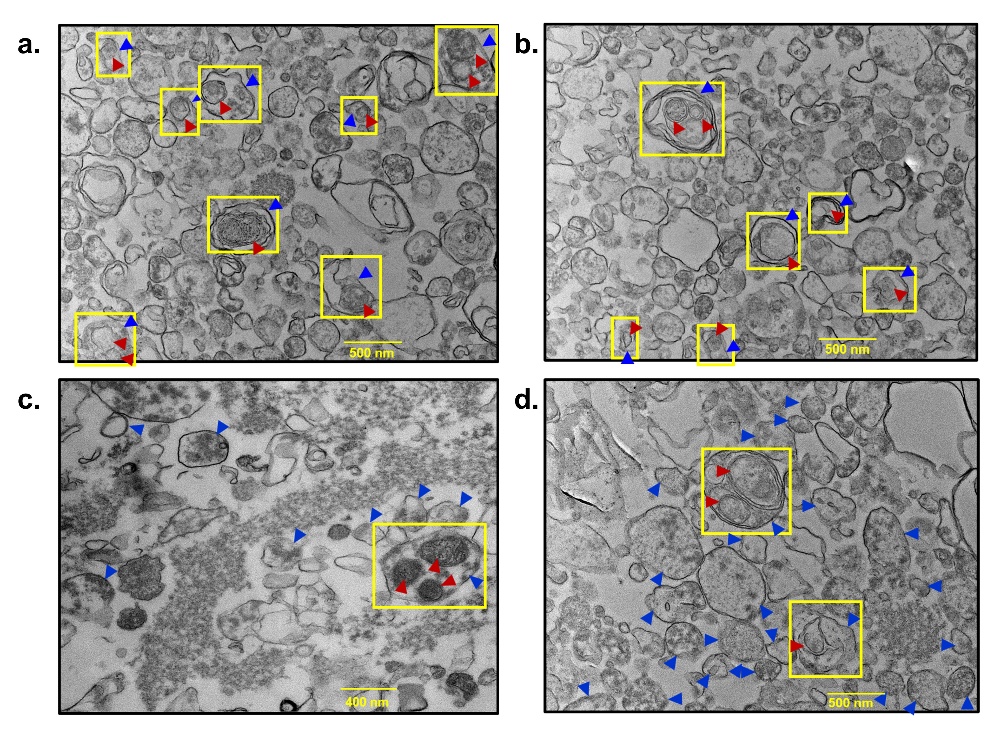

**Supplemental Figure 2: Representative TEM images used for quantification of mitochondrial incorporation within SC-lEVs.** Analyses included the proportion of mitochondria-containing and mitochondria-negative lEVs, as well as the distribution of vesicles containing one or multiple mitochondria. Yellow boxes denote representative mitochondria-containing lEVs. Blue arrowheads indicate lEVs, whereas maroon arrowheads indicate mitochondria enclosed within lEVs. Scale bars = 400–500 nm.

**Supplemental Table 1: Quantification of calcein-positive EVs and calcein/MitoTracker double-positive mitochondria-containing extracellular vesicles within the total particle population of SC-lEVs and SC-sEVs using ImageStream flow cytometry.**

|  | **Total EV events** | **Calcein-positive events** | **Calcein-positive EVs (%)** | **Calcein-AM/MitoTracker double-positive events** | **Calcein-AM/MitoTracker double-positive EVs (%)** |
| --- | --- | --- | --- | --- | --- |
| **SC-lEVs** | 81,657 | 53,971 | **66.1** | 446 | **0.83** |
| **SC-sEVs** | 1,32,937 | 93,774 | **70.5** | 37 | **0.04** |

**Supplemental Table 2: Quantitative assessment of mitochondrial incorporation within SC-lEVs, including the percentage of mitochondria-containing lEVs, mitochondria-negative lEVs, and the distribution of vesicles harboring one or multiple mitochondria.**

| **TEM image frame number** | **Total SC-lEVs** | **Mitochondria-containing SC-lEVs** | **Empty SC-lEVs** | **Mitochondria-containing SC-lEVs (%)** | **Empty SC-lEVs (%)** | **Total number of SC-lEVs having one mitochondrion** | **Total number of SC-lEVs having two mitochondria** | **Total number of SC-lEVs having three mitochondria** |
| --- | --- | --- | --- | --- | --- | --- | --- | --- |
| 1 | 58 | 8 | 50 | 13.8 | 86.2 | 12 (70.6%) | 4 (23.5%) | 1 (5.9%) |
| 2 | 66 | 6 | 60 | 9.1 | 90.9 |  |  |  |
| 3 | 8 | 1 | 7 | 12.5 | 87.5 |  |  |  |
| 4 | 25 | 2 | 23 | 8.0 | 92.0 |  |  |  |

**Supplemental Table 3: Measured densities of fractions collected from the iodixanol density gradient during lEV purification.**

| **EVs used for the study** | **Fractions** | **Average Density (g/mL)** |
| --- | --- | --- |
| **DGC-lEVs** | 1 | 1.027±0.004 |
|  | 2 | 1.034±0.006 |
|  | 3 | 1.041±0.004 |
|  | 4 | 1.061±0.003 |
|  | 5 | 1.078±0.003 |
|  | 6 | 1.093±0.007 |
|  | 7 | 1.102±0.021 |
| **NVEP-lEVs** | 8 | 1.170±0.010 |
|  | 9 | 1.185±0.006 |
|  | 10 | 1.217±0.027 |
|  | 11 | 1.204±0.004 |
|  | 12 | not available |
